# Rapid green leaf volatile sensing allows intact plants to release induced volatiles earlier than wounded emitters

**DOI:** 10.64898/2026.08.24.746686

**Authors:** Tristan M. Cofer, Lingfei Hu, Gaetan Glauser, Matthias Erb

## Abstract

Organisms in the vicinity of attacked neighbors can respond to danger cues by activating their own defenses. However, the temporal dynamics of these interactions remain poorly resolved, despite their importance for the ensuing effects. Here, we investigated defense responses of maize plants that are exposed to induced volatiles from herbivory-induced neighbors in real time. These experiments revealed a surprising temporal paradox: Intact neighbors start emitting most induced volatiles 15–80 min earlier than the attacked plants themselves. This earlier responsiveness was not associated with earlier induction of defense hormones, volatile biosynthesis genes or within-leaf volatile accumulation. Instead, receiver plants kept their stomata open, while attacked sender plants rapidly closed them to combat water loss. Using different volatile deficient mutants, we demonstrate that the rapid responses in receivers are triggered by green leaf volatiles, which are released from the wound sites of sender plants within minutes independently of stomatal aperture. Thus, non-attacked neighbors can act as “unhindered observers” that activate some defenses more quickly than attacked organisms themselves, as they do not have to deal with wound trauma. The resulting plant volatile concatenation patterns expand our understanding of volatile signaling and have the potential to shape population-level multitrophic interactions.

## Introduction

The capacity to use early warning cues to anticipate incoming attacks is essential for effective defense and immunity. Plants can use a series of chemical cues to detect herbivory before the onset of attack. These cues include insect pheromones (1), egg-derived cues (2, 3), and volatiles from attacked neighboring plants (4–6). Volatile-mediated defense induction is common across many different plant species (7, 8) and can protect plants from herbivory in natural and agricultural settings (9–14). Yet, we often lack a detailed picture of the underlying temporal dynamics, thus constraining our understanding of these interactions. Timing is likely a key determinant of volatile-mediated interactions between plants, herbivores and their natural enemies, as it affects herbivore host plant choice and predator foraging success (15, 16), and thus warrants a detailed understanding.

Different types of induced volatiles can enhance defense responses in non-attacked tissues and neighboring plants (7). The growing list includes homo-and monoterpenes (4, 6, 17, 18), aromatic compounds (2, 19, 20), jasmonates (14, 21, 22) and green leaf volatiles (5, 23–25). Green leaf volatiles are released rapidly upon wounding and can trigger defense responses in a variety of different plants (26, 27). In maize, green leaf volatiles that are emitted by attacked neighbors have been found to be essential for the induction of systemic root responses (12), while indole from attacked neighbors has been associated with leaf defense priming (19). Experiments with synthetic green leaf volatiles show that they can directly induce and prime maize defenses (28, 29).

Volatile exposure results in defense activation via several distinct physiological steps (30, 31). First, volatiles enter the leaves, primarily through the stomata (32). Second, the volatiles are perceived through (as yet unknown) receptors (29, 33). Third, following early signaling events (34–36), defense hormones are deployed. In many cases, herbivory-induced volatiles induce and/or prime jasmonate signaling through the accumulation of jasmonic acid (JA) and its isoleucine conjugate (JA-Ile) (12, 28, 37, 38). Fourth, jasmonate signaling triggers the expression of defense genes, including biosynthesis genes of defensive metabolites, leading to the accumulation of defense compounds (25).

Interestingly, some plants start producing volatiles themselves upon stress volatile exposure (4, 39). In this case, the volatiles are produced *de novo* following the induction of their biosynthesis genes (4, 40, 41), transported out of the cells (42) and likely released through the stomata (43). These induced volatiles can then again enhance plant resistance, attract natural enemies, repel or attract herbivores and potentially trigger further responses in other plants (4, 44). Thereby, they can have important consequences for multitrophic interaction networks (15).

The speed and timing of defense induction in sender and receiver plants likely plays a pivotal role in conveying protection for plants (45). Green leaf volatiles are triggered within seconds to minutes, while terpenes and aromatic compounds are released within hours following attack (46, 47), thus providing different temporal windows for stress detection by neighbors. Studies show that direct neighbor responses can be induced relatively quickly, with plants responding to green leaf volatiles within minutes (34, 35). Defense priming is typically observed at later timepoints, with studies often employing delays of several hours to days after stress volatile exposure to measure the intensity of induction upon elicitation by a separate stressor (19, 48). In maize, first volatile responses in receiver plants can be detected within hours after the start of caterpillar attack of sender plants, followed by a second “primed” volatile burst after day two (39). Yet, despite many years of research, the exact timing of the volatile exchange between sender and receiver plants is unknown. What is the time lag between the attack of a sender plant and the defense response, including volatile release, of a neighboring plant? This question remains unresolved, despite its potential importance.

Here, we took advantage of the fact that herbivory induced sender and non-attacked receiver maize plants respond by releasing herbivore-induced plant volatiles such as indole and various terpenes (39) to compare defense activation in both plants simultaneously in real time. Surprisingly, we uncovered a temporal paradox: receiver plants released induced volatiles more quickly than sender plants. To explain this phenomenon, we measured the speed of upstream and regulatory responses, including jasmonate accumulation, volatile biosynthesis gene expression, within-leaf accumulation of volatiles, and stomatal aperture. We then generated and used different volatile biosynthesis mutants as senders to identify the early released sender volatiles that resolve the apparent paradox and allow receiver plants to respond more quickly.

## Results

### Receiver plants emit most volatiles more rapidly than herbivore-induced sender plants

To determine the speed at which plants respond to volatiles from attacked neighbors, we conducted a detailed time-course analysis of volatile induction in sender and receiver plants. Sender plants were induced by wounding and application of insect oral secretions three times, at 0, 30, and 60 min after the start of the experiment. Volatiles were then measured every 15 min using a custom robotic high-throughput proton transfer reaction time-of-flight mass spectrometry (PTR-ToF-MS) system (39, 49). Within minutes after simulated herbivory, sender plants released green leaf volatiles, detected in our experiment as m/z 99.08, the protonated mass of (*Z*)-3-and (*E*)-2-hexenal (C_6_H_11_O^+^) (Fig. 1). Receivers did not emit hexenal, but absorbed a portion of hexenal emitted by the sender plants (Fig. 1). After one to two hours, both sender and receiver plants started to release indole, monoterpenes, sesquiterpenes, and the homoterpenes (*E*)-4,8-dimethyl-1,3,7-nonatriene (DMNT) and (3*E*,7*E*)-4,8,12-trimethyltrideca-1,3,7,11-tetraene (TMTT; Fig. 1). Surprisingly, the emissions of all these volatiles started 15–80 min earlier in receiver than in sender plants. Their emissions also peaked 30 min to 4 h earlier in receivers than in senders. For indole, DMNT and TMTT, we also detected total higher emissions from receiver than from sender plants. To verify the surprising earlier release by receiver plants with an orthogonal method, we collected volatiles using a traditional push-pull system (46) and quantified them by gas chromatography / mass spectrometry (GC/MS). We specifically collected and quantified volatiles from 60 to 75 min, which falls within the window of earlier release of some of the volatiles in receiver plants. During this time window, we detected indole and linalool emissions from receiver plants, but not yet from sender plants (Fig. S1). DMNT was induced in both sender and receiver plants, with higher levels in receiver plants. TMTT and sesquiterpenes were not yet detected. To evaluate the impact of early volatile exposure on emission speed, we performed an experiment where receivers were disconnected from the senders 75 min after sender treatment, and senders and receiver volatiles were measured separately by PTR-ToF-MS. Receiver plants emitted indole and terpenes earlier than receivers, similar to the first experiment (Fig. S2). Finally, we pulse-labelled receiver plants with ^13^CO_2_ to specifically test for the release of *de novo* synthesized volatiles in receivers. We achieved significant incorporation of ^13^C into DMNT (Fig. S3), allowing us to quantify [¹³C_1_]DMNT and [¹³C_2_]DMNT dynamics in labeled receivers by PTR-ToF-MS. Labeled DMNT was released earlier from labeled receivers than unlabeled DMNT from unlabeled senders (Fig. S3). Thus, non-attacked receiver plants consistently release induced volatiles (other than GLVs) earlier than herbivory-induced sender plants.

**Figure 1.**
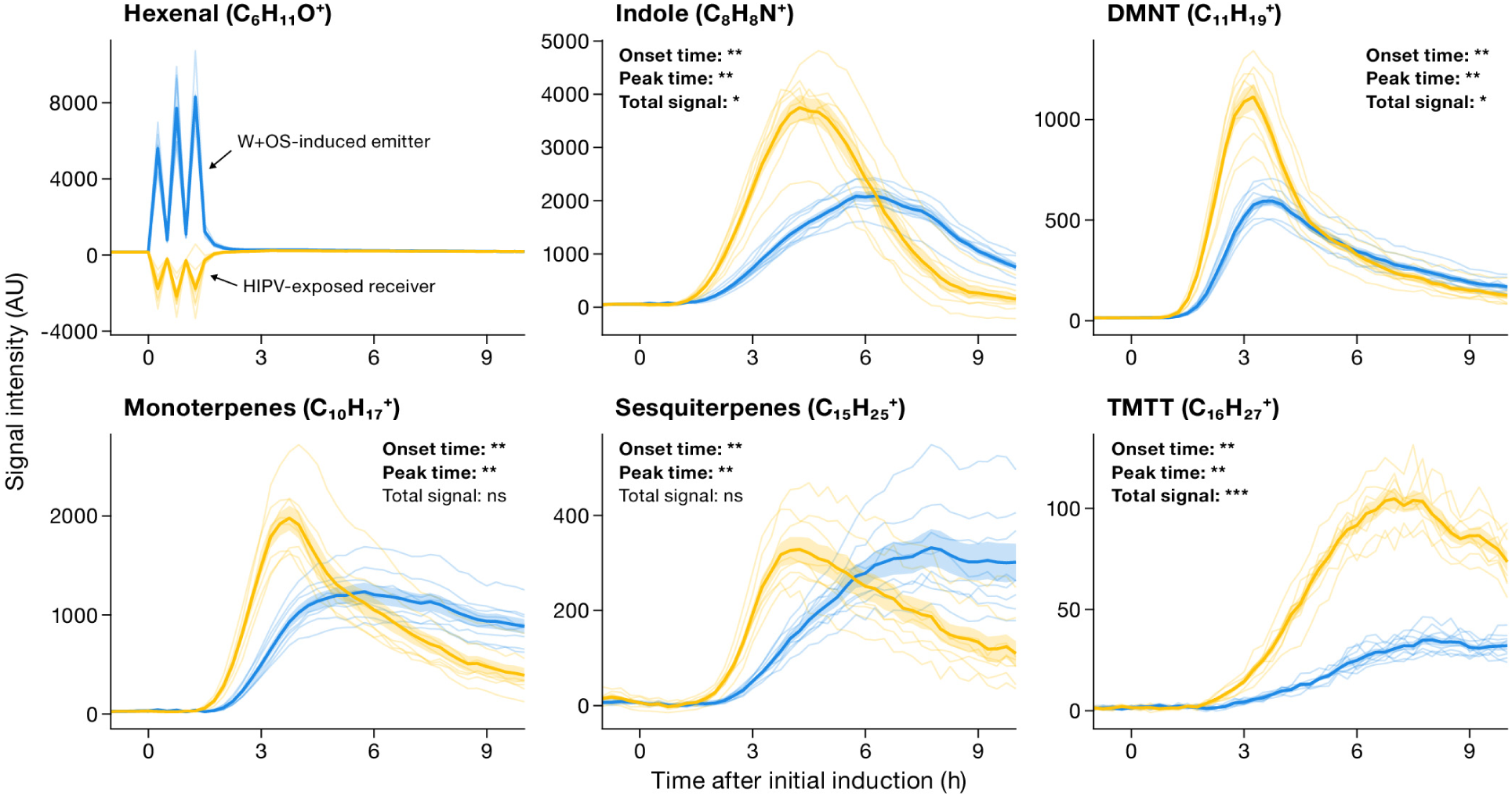
Undamaged receiver plants emit most stress-related volatiles earlier than herbivory-induced sender plants. Sender plants were induced by mechanical wounding followed by application of insect oral secretions at 0, 30, and 60 min. Volatile emissions were monitored every 15 min from sender plants alone and from sender plants paired with undamaged receiver plants using a custom-built, robotic high-throughput proton transfer reaction time-of-flight mass spectrometry (PTR-ToF-MS) system. Emissions from receiver plants were calculated by subtracting emissions measured from sender plants alone from those measured in sender-receiver pairs. Stress-related volatiles, including (*Z*)-3-and (*E*)-2-hexenal (combined signal), indole, mono-, sesqui-, and homoterpenes (DMNT, TMTT), were detected as protonated ions based on characteristic *m/z* signals. Solid lines represent mean emission rates for sender (blue) and receiver (yellow) plants; shaded ribbons indicate ± SEM, and thin lines show individual biological replicates (n = 8). For each analyte, onset time, peak time, and cumulative signal intensity were quantified from baseline-normalized time courses. Onset time was estimated by change-point analysis, peak time was defined as the earliest time point of maximum signal, and cumulative total signal intensity was calculated as the trapezoidal area under the time course. Statistical differences between sender and receiver plants were assessed using two-sided Wilcoxon rank-sum tests with Benjamini-Hochberg correction for multiple comparisons. Asterisks indicate significance levels (\**P* < 0.05; \*\**P* < 0.01; *\*\*\*P* < 0.001); ns, not significant.

### Earlier HIPV release is not associated with stronger Hormonal or Transcriptional responses in Receiver plants

Plant volatile emissions are regulated by phytohormone signaling and transcriptional activation of biosynthetic genes (50, 51). To investigate whether differences in these regulatory steps can explain the more rapid volatile emissions of receiver plants, we measured the induction of phytohormones and key volatile biosynthesis genes in a detailed time series (Fig. 2A). Jasmonic acid (JA), JA-isoleucine (JA-Ile), and abscisic acid (ABA) were induced in both sender and receiver plants within 15–45 min, with similar induction kinetics and overall higher levels in sender plants. 12-oxo-phytodienoic acid (OPDA) was induced significantly in sender plants after 15 min, but remained unchanged in receivers. Salicylic acid (SA) concentrations were induced at later time points in receiver, but not sender plants. Thus, the more rapid volatile emissions in receiver plants are not associated with a quicker or stronger phytohormone burst. Transcript levels of the terpene biosynthesis genes *FPPS3*, *TPS2*, *TPS3*, *TPS10*, and *CYP92C5* (51, 52), as well as the indole biosynthesis gene *IGL* (40), were significantly upregulated after 15–75 min in both sender and receiver plants compared to controls (Fig. 2B). Induction speed was similar in sender and receiver plants, with higher transcript levels of all genes apart from *IGL* at later timepoints in sender plants. Thus, the more rapid volatile emissions in receiver plants are not associated with a quicker transcriptional activation of the underlying biosynthesis genes.

**Figure 2.**
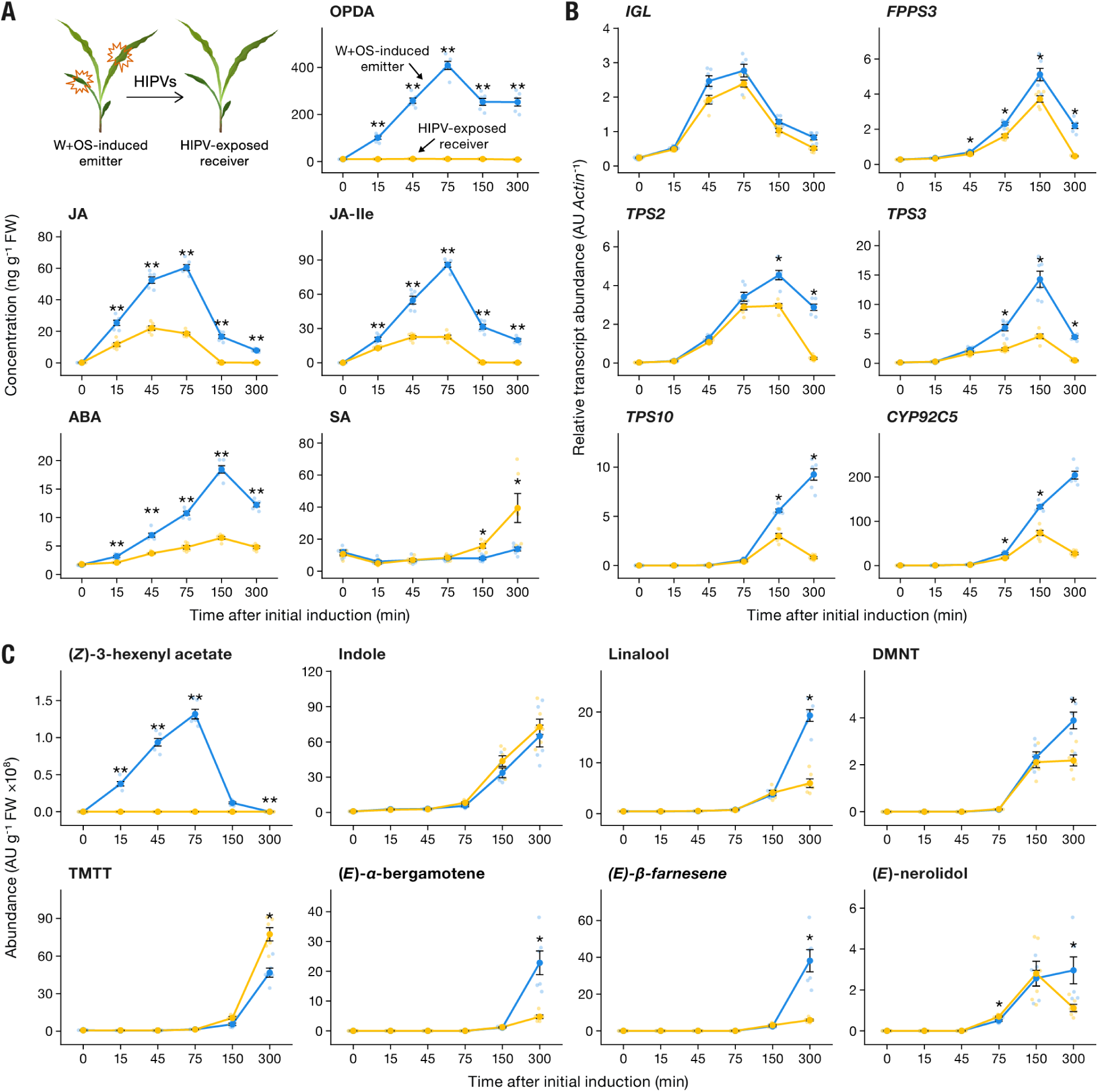
Early volatile release in receiver plants is not associated with stronger hormonal or transcriptional responses. Sender plants were induced by mechanical wounding followed by application of insect oral secretions at 0, 30, and 60 min. The schematic in panel A illustrates the W+OS-induced sender plant and the HIPV-exposed receiver plant. (A) Stress-related phytohormones (OPDA, JA, JA-Ile, ABA, and SA), (B) transcript levels of volatile biosynthesis genes (*IGL*, *FPPS3*, *TPS2*, *TPS3*, *TPS10*, and *CYP92C5*), and (C) free within-leaf pools of volatile compounds, including (*Z*)-3-hexenyl acetate, indole, linalool, DMNT, TMTT, (*E*)-*α*-bergamotene, (*E*)-*β*-farnesene, and (*E*)-nerolidol, were quantified in the same sender (blue) and receiver (yellow) plants at the indicated time points. Points represent individual biological replicates; lines connect mean values; error bars indicate ± SEM (n = 6). At each time point, sender and receiver values were compared using two-sided Wilcoxon rank-sum tests with Benjamini-Hochberg correction for multiple comparisons within each analyte. Asterisks indicate significance (\**P* < 0.05; \*\**P* < 0.01); absence of asterisks indicates no significant difference.

To investigate whether receiver leaves accumulate volatiles earlier than sender leaves, we quantified free within-leaf pools of (*Z*)-3-hexenyl acetate, indole, linalool, DMNT, TMTT, (*E*)-*α*-bergamotene, (*E*)-*β*-farnesene, and (*E*)-nerolidol at different time points (Fig. 2C). (*Z*)-3-Hexenyl acetate rapidly accumulated in sender plants upon simulated herbivory, but not in receiver plants. The other volatiles accumulated in the leaves of both sender and receiver plants 75 to 150 min after elicitation of the sender plants. Accumulation kinetics were overall similar, with linalool, DMNT, (*E*)-*α*-bergamotene and (*E*)-*β*-farnesene accumulation being higher in sender plants 300 min after elicitation. Thus, the more rapid emission of volatiles in receiver plants is not associated with a more rapid accumulation in the leaves. Together, these experiments demonstrate that neither hormonal defense signaling nor volatile biosynthesis *per se* can explain the more rapid release of volatiles from receiver plants.

### Receiver plants escape wound-induced Stomatal Closure

Wounding leads to stomatal closure to combat water loss, which may impair the release of volatiles (53, 54). We hypothesized that receiver plants may be less constrained by stomatal closure, which may allow them to release volatiles more rapidly. Indeed, sender plants started closing the stomata of the herbivory-induced leaves within 15 min, resulting in a reduction of stomatal conductance of more than 50% after 45 min (Fig. 3A). By contrast, receiver plants kept their stomata open, and only showed a mild reduction at 150 min after sender exposure. To determine whether the reduction in stomatal conductance observed in sender plants is triggered by herbivore-specific cues, we wounded leaves and applied either insect oral secretions or water (Fig. 3B). The decrease in stomatal conductance was similar between treatments, showing that this is a wound-induced response. To determine whether stomatal closure occurs systemically in sender plants, we measured stomatal conductance in the second oldest leaf when either this leaf was herbivory-induced (local treatment) or the third oldest leaf was herbivory-induced (systemic treatment). Stomatal conductance was reduced only when the measured leaf itself was herbivory-induced (Fig. 3C). Thus, herbivory-induced leaves of sender plants close their stomata rapidly, while receiver plants show a much weaker response.

**Figure 3.**
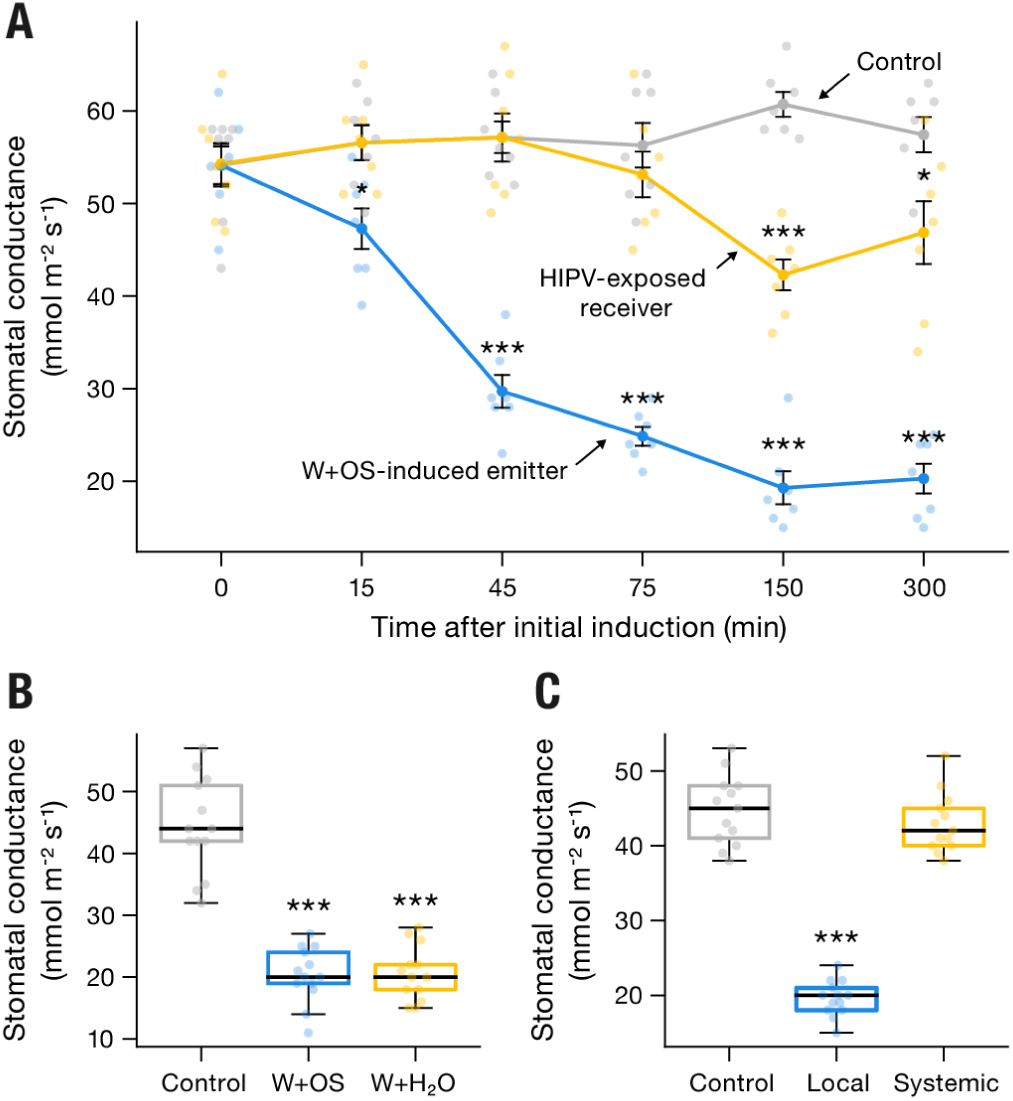
Sender plants rapidly reduce stomatal conductance, whereas receiver plants maintain open stomata. (A) Sender plants were induced by mechanical wounding followed by application of insect oral secretions at 0, 30, and 60 min, and stomatal conductance was measured over time in the second oldest leaf of sender plants (blue), receiver plants (yellow), and untreated control plants (gray). Points represent individual biological replicates, and lines indicate mean values; error bars indicate ± SEM (n = 7) (B) Stomatal conductance measured 75 min after induction in wounded leaves treated with insect oral secretions or water. (C) Stomatal conductance was measured 75 min after local induction of the second oldest leaf or after systemic induction of the third oldest leaf. Box plots show medians, interquartile ranges, and individual data points (n = 13). Differences among treatments were analyzed using one-way ANOVA followed by Tukey’s honestly significant difference (HSD) test for post hoc comparisons. Asterisks indicate significant differences relative to the control (\**P* < 0.05; \*\**P* < 0.01; *\*\*\*P* < 0.001); absence of asterisks indicates no significant difference.

To better understand the role of stomata in conveying earlier neighbor responses, we thought to manipulate their behavior in sender and receiver plants. Opening stomata with the fungal toxin fusicoccin (55) or by genetically manipulating the anion channel SLAC1 (56) led to pleiotropic effects on plant growth and health, and was thus not informative to measure the speed of volatile release from wounded sender plants. Closing stomata of receiver plants by ABA treatment on the other hand restricts the capacity of the receivers to take up and respond to the volatiles of sender plants (57), and was thus equally unpractical. Thus, stomatal effects are associated with the earlier neighbor responses, but whether they are causal cannot be determined with currently available experimental approaches.

### Green Leaf Volatiles Are required for Rapid Responses in Neighboring Plants

To uncover which volatiles are responsible for triggering rapid neighbor responses, we determined volatile release of wild type receiver plants to herbivory-induced volatiles from different mutant sender plants with defects in the major volatile biosynthesis pathways. First, we used *igl* mutants, which are deficient in indole biosynthesis (19, 40). As expected, *igl* mutant plants did not emit any indole upon elicitation (Fig. S4). Hexenal levels were similar to wild type plants. The other volatiles were emitted at slightly lower levels in the mutant plants. Receiver plants displayed similar hexenal uptake and volatile induction kinetics upon exposure to *igl* mutants compared to wild type plants (Fig. S4), showing that indole is not required for this rapid direct response. Second, we generated a *tps2/3* double knockout, expected to be defective in mono-and homoterpene biosynthesis (58). *tps2/3* mutant sender plants showed a significantly lower induction of monoterpenes, DMNT and TMTT, but similar emissions of hexenal (Fig. S4). Indole and sesquiterpene emissions were slightly reduced compared with wild type plants. Receiver plants displayed similar volatile induction kinetics upon exposure to induced *tps2/3* mutants compared to wild type plants (Fig. S4). Thus, mono-and homoterpenes are not required for the rapid response of receiver plants. Third, we created a *lox10* mutant that is defective in green leaf volatile biosynthesis (59). As expected, the *lox10* sender plants showed non induction of hexenal upon elicitation, and also emitted significantly lower levels of all other induced volatiles (Fig. 4), as reported before for *lox10* mutants in B73 (59). Remarkably, receiver plants completely failed to respond to herbivory-induced *lox10* plants. For both indole and terpenes, we even found negative emission rates, suggesting net uptake by the receivers in the absence of induction by green leaf volatiles. (Fig. 4). Thus, green leaf volatiles appear to be necessary to trigger rapid responses and volatile emissions, which is in line with their early emission and the fact that early volatiles (emitted before 75 min) trigger rapid responses in receivers (Fig. S2).

**Figure 4.**
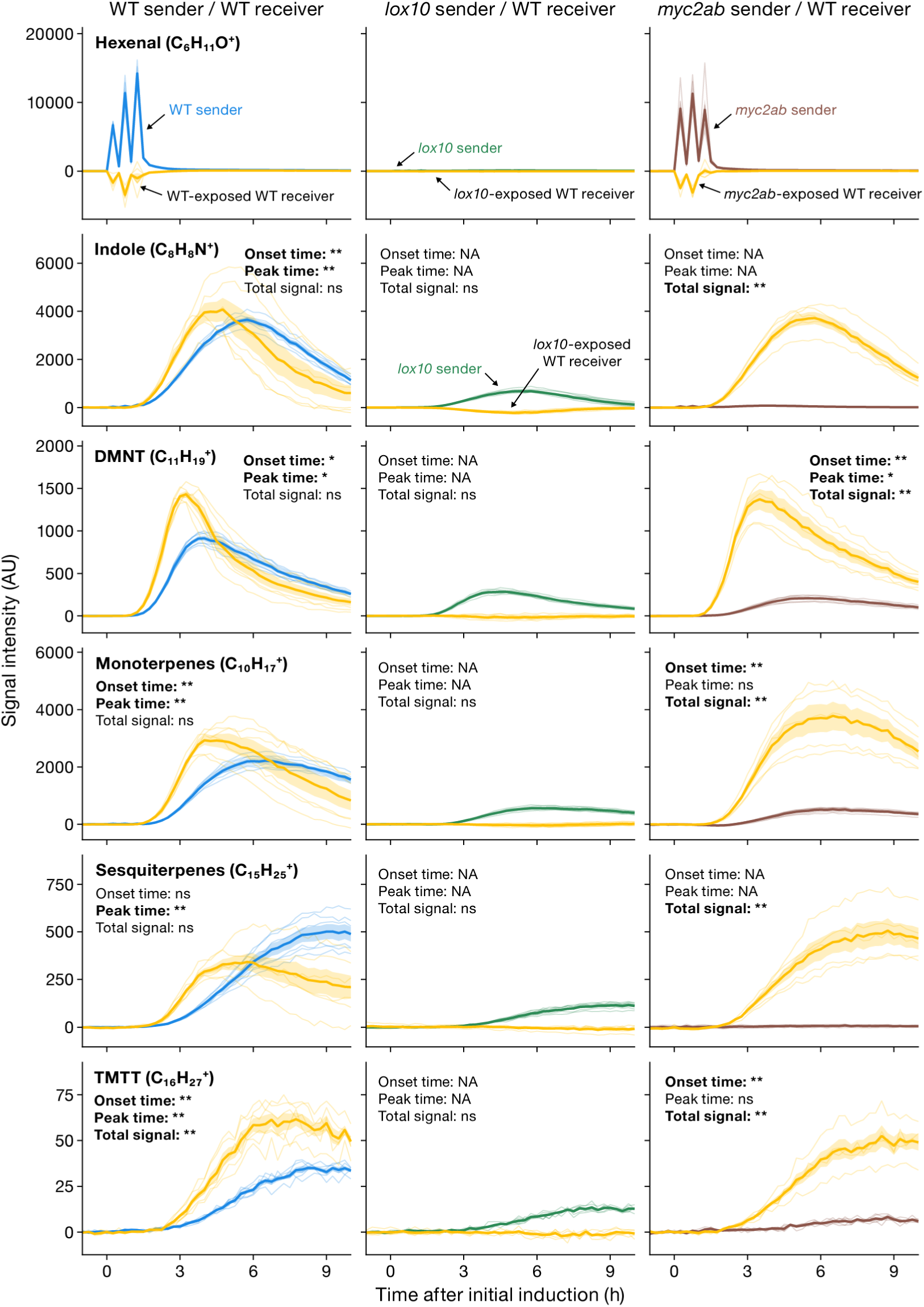
Green leaf volatiles are required for rapid responses in neighboring plants. Time-resolved volatile emissions were measured from wild type (WT) receiver plants exposed to herbivory-induced WT or mutant sender plants that are unable to emit certain volatiles. Sender plants were induced by mechanical wounding followed by application of insect oral secretions at 0, 30, and 60 min, and volatile emissions were monitored every 15 min from sender plants alone and from sender-receiver pairs. Receiver emissions were calculated by subtracting emissions from sender plants alone from those measured in sender-receiver pairs. Shown are emissions from WT, *lox10* and *myc2ab* sender plants paired with WT receivers. The *lox10* mutant is deficient in green leaf volatile biosynthesis, whereas *myc2ab* is impaired in jasmonate signaling and induction of most herbivore-induced volatiles but retains green leaf volatile emissions. Stress-related volatiles, including (*Z*)-3-and (*E*)-2-hexenal (combined signal), indole, mono-, sesqui-, and homoterpenes (DMNT, TMTT), were detected as protonated ions based on characteristic *m/z* signals. Solid lines represent mean emission rates for sender plants (blue, green, or brown) and WT receiver plants (yellow); shaded ribbons indicate ± SEM, and thin lines show individual biological replicates (n = 6). For each analyte, onset time, peak time, and cumulative signal intensity were quantified from baseline-normalized time courses. Statistical differences were assessed using two-sided Wilcoxon rank-sum tests with Benjamini-Hochberg correction for multiple comparisons. Asterisks indicate significance levels (\**P* < 0.05; \*\**P* < 0.01); ns, not significant; NA, metric not determined. For other volatile mutants, see Fig. S4.

As the *lox10* sender plants are emitting lower levels of other induced volatiles, we sought to further confirm the role of green leaf volatiles. To do so, we employed *myc2ab* mutants as sender plants. This mutant is defective in jasmonate signaling and the induction of volatile biosynthesis genes (50). We found that *myc2ab* mutants emit no to very low levels of all induced volatiles, apart from green leaf volatiles, whose emissions were fully intact (Fig. 4). Receiver plants showed fully intact rapid responses when exposed to induced volatiles of *myc2ab* mutants (Fig. 4). This result confirms that none of the other induced volatiles apart from green leaf volatiles are required for the rapid induction of neighboring plants.

## Discussion

Induced responses across populations are shaped by a dynamic interplay of direct responses to stressors and indirect responses to individuals reacting to these stressors. From fish swarms attacked by predators (60) to microbial communities invaded by bacteriovores (61), bees sacrificing themselves to protect the hive (62), and humans working together to stop the spread of contagion (63), this interplay determines population-level behavior and interaction outcomes. The time lag in defense activation between attacked individuals and non-attacked neighbors is a critical, yet poorly resolved component of these situations. This is particularly true for volatile interactions between plants, where limited temporal resolution restricts our ability to capture mechanisms and ecological consequences.

Plants can respond relatively quickly to volatiles from stressed neighbors (39). Membrane depolarization, cytosolic calcium influx, and MPK phosphorylation occur within seconds to minutes in plants that are exposed to these volatiles (34–36). Given that i) the first stress volatiles from attacked plants peak after 0.5 minutes (47) and that they are rapidly transmitted to neighbors that are either downwind or in close proximity (10), we expected the lag time of terpene and indole release responses in maize to be less than 1 minute compared to the sender responses. To our surprise, we found that there was no lag time; on the contrary, neighbors released most induced volatiles 15–80 min earlier than the senders. This discovery demonstrates that non-attacked plants show distinct and unexpected induction kinetics and can activate some of their volatile defenses more effectively than attacked plants.

What is the mechanism behind the more rapid volatile release of non-attacked plants? We tested the hypothesis that the defense-inducing volatiles may trigger defense signaling more quickly than the non-volatile elicitors at the wound site. Wounded plants need to balance defense and regeneration, which are subject to signaling trade-offs (64, 65) that may constrain the speed of defense induction. Based on our data, we can reject the hypothesis that sender plants are slower in defense signaling deployment *per se*, as both sender and receiver plants deploy defense hormones, volatile biosynthesis genes, and volatile production at a similar speed. Thus, the most parsimonious explanation is that receiver plants must be able to release their induced volatiles more quickly. Wounding leads to rapid water loss, thus triggering stomatal closure (53, 66). We demonstrate that this response is largely restricted to wounded sender leaves and does not occur in unwounded receivers at the same level. Several studies show a strong link between stomatal aperture and the release of terpenes and indole (54, 67). Stomatal closure is likely strongest close to the wound site. As intact cells around the wound site also produce the majority of the wound-induced volatiles (68, 69), the volatiles have to diffuse through leaf air spaces to reach wounds or open stomata further away from the wound site, thus likely delaying their release. Complete stomatal closure around the wound site and diffusion through air spaces could explain why the release of volatiles with a high Henry’s Law Constant such as homoterpenes and sesquiterpenes is also delayed, despite the fact that their release should not be affected strongly by reduced stomatal conductance (70). An alternative hypothesis is that volatile transport processes into the apoplast are delayed in wounded plants compared to receiver plants. A better understanding of the mechanisms of volatile transport and release (71) would be necessary to distinguish these hypotheses.

Which volatiles are driving the rapid volatile release of neighboring plants? Green leaf volatiles are well known for their capacity to directly induce defenses, including volatiles (27). The release of (*Z*)-3-hexenal and (*Z*)-3-hexen-1-ol happens quickly upon wounding and is mostly unconstrained by stomatal aperture (72), thus making these volatiles prime candidates to trigger rapid volatile release in neighbors. Using a combination of volatile biosynthesis mutants, we show that green leaf volatiles are required for triggering the rapid neighbor responses, in line with their rapid, stomata-independent release from senders. While green leaf volatiles are required for direct defense induction, other volatiles such as indole, which are induced rapidly by green leaf volatiles in receivers, can prime defenses alone and in combination with green leaf volatiles at later timepoints (19, 73), but are not required for the rapid neighbor response. How plants integrate multiple active volatiles during their rapid and dynamic volatile mediated interactions is an interesting avenue for future research.

The biological consequences of the more rapid volatile release from non-damaged neighbors are potentially significant. Here, we discuss two scenarios by which this phenomenon could influence multitrophic interactions. First, by rapidly releasing volatiles that can prime and act in synergy with green leaf volatiles to regulate defenses, the neighbors may be part of a feedback loop that strengthens defense expression of the sender plant. Similar feedbacks could also happen between attacked and non-attacked leaves of the same plants (11, 25), thus resulting in rapid systemic induced resistance patterns. Second, the neighbor could engage in a form of relay cueing to rapidly enhance the distance an intensity of “alarm calls” which may then boost natural enemy attraction to the attacked plants. It would be interesting to explore if predators and parasitoids can use induced volatiles for long range location of infested patches, and green leaf volatiles for short range host location (74), in which case rapidly responding neighbors could serve as guideposts for effective biological control. From an adaptive point of view, these scenarios could represent a form of manipulation, where the sender plant uses the receiver to strengthen defense signaling. Or they could be a form of cooperation, where plants interact to augment each other’s defenses (9, 75). The latter would be most likely for plants that are closely related, including clonal plants. In the case of maize, it is interesting to note that its wild ancestor teosinte branches heavily – the patterns observed in cultivated, genetically identical plants may thus be a remnant of within plant, branch-to-branch communication (11). Further experiments will be necessary to test these hypotheses. Higher order mutants that are defective in several induced volatiles as well as mutants that are deficient in volatile perception could be useful tools to explore to what extend “unhindered neighbors” influence multitrophic interactions in natural and agricultural systems.

## Materials and Methods

### Plant material and growth conditions

The *lox10* and *tps2/3* knockout lines were generated in the maize inbred line KN5585 using CRISPR/Cas9 technology (Figure S5 and S6). The *igl* and *myc2ab* mutants in the KN5585 background have been described (12, 50). Maize plants were grown in commercial potting soil (Selmaterra; Bigler AG, Steffisburg, Switzerland) in a climate-controlled greenhouse at 22 ± 2°C, 40–60% relative humidity, and a 14:10 h light/dark cycle with supplemental lighting (∼300 μmol m⁻² s⁻¹). Plants at the V2 stage, with three fully expanded leaves, were used for all experiments.

### Experimental setup

Individual sender plants were enclosed in cylindrical glass chambers (12 cm diameter × 45 cm height), each connected via Teflon tubing to a second identical chamber that either remained empty or contained a single receiver plant. Clean air was supplied to the sender chamber at 0.8 L min⁻¹ and exited through the receiver chamber, allowing volatiles to pass from sender to receiver. LED lights (DYNA, Heliospectra), positioned ∼80 cm above the chambers, provided illumination at ∼300 μmol m⁻² s⁻¹. The light/dark cycle followed the same 14:10 h schedule as used in the greenhouse.

### Plant induction by simulated herbivory

Sender plants were induced by wounding leaves 2 and 3 on both sides of the central vein over an area of 0.5 cm² using a fine metal grater. Each wound site was then treated with 5 μL of *Spodoptera exigua* oral secretions diluted 1:1 in Milli-Q water. To mimic continuous feeding, the induction treatment was repeated three times at 30-minute intervals, sequentially targeting the base, middle, and tip of each leaf.

### Receiver-disconnection experiment

Sender and receiver plants were connected, and sender plants were induced as described above. After 75 min, receiver plants were disconnected from the sender chambers and supplied individually with clean air at the same flow rate. Sender and receiver plants were then monitored for volatile release separately by PTR-ToF-MS for the remainder of the experiment as described below.

### Real-time volatile monitoring

Volatile emissions were monitored in real time using a modified proton transfer reaction time-of-flight mass spectrometer (PTR-ToF-MS; TOFWERK, Switzerland) coupled with a custom-made automated headspace sampling system (Abon Life Sciences, Switzerland). The receiver chamber outlet was connected to the PTR-ToF-MS, which continuously sampled air at 0.1 L min⁻¹. Each chamber was sampled every 15 min, with volatiles recorded continuously for 15–25 s per time point and summarized as the median value. A zero-gas flush was included between samples to prevent carryover. Mass spectra (0–500 m/z) were acquired in positive ion mode at ∼10 Hz, with the PTR operated at 100°C and an E/N ratio of ∼120 Td. Data were analyzed using Tofware v3.2.2, and compounds were identified based on their protonated molecular masses ([M+1] *m/z*; Fig. S7).

### Volatile collection and analysis by GC-MS

To complement real-time PTR-ToF-MS analysis, volatiles were collected for gas chromatography-mass spectrometry (GC-MS) using a push-pull headspace sampling system. Clean air was pushed into the sender chamber and simultaneously pulled from the receiver chamber at equal flow rates of 0.8 L min⁻¹. Volatiles were trapped on SuperQ adsorbent filters (20 mg; Alltech Associates, Deerfield, IL, USA) over a defined interval (e.g., 60–75 min after the first induction treatment). Filters were eluted with 100 µl of hexane/dichloromethane (1:1, v/v), and samples were analyzed using an Agilent 7820A gas chromatograph coupled to an Agilent 5977E quadrupole mass selective detector. The MS was operated with an interface temperature of 280°C, quadrupole temperature of 150°C, source temperature of 230°C, and electron energy of 70 eV. Sample aliquots were injected in splitless mode onto an HP-5MS capillary column (30 m × 0.25 mm i.d. × 0.25 μm film thickness; Agilent, Palo Alto, CA, USA) with helium as the carrier gas at a constant flow rate of 1 mL min⁻¹. The oven temperature was held at 40°C for 4 min, then ramped to 200°C at 5°C min⁻¹. Volatile compounds were identified by comparison with the NIST Mass Spectral Library (USA) and, when available, with authentic standards, and quantified based on the integrated areas of individual chromatographic peaks (Fig. S8).

### ^13^CO_2_ PULSE-LABELING EXPERIMENT

In a preliminary experiment, plants were pulse-labeled with ^13^CO_2_ or ^12^CO_2_ gas (Sigma-Aldrich) to determine whether ^13^C was incorporated into herbivore-induced volatiles. Thirty minutes before the onset of the light period, individual plants were transferred to sealed 5-L glass chambers with the inlet and outlet ports closed. A 5-mL aliquot of ^13^CO_2_ or ^12^CO_2_ gas (Sigma-Aldrich) was injected through an inlet fitted with a Teflon septum using a gas-tight syringe, yielding a nominal initial chamber concentration of approximately 1,000 ppm (v/v). After 1.5 h of illumination, an additional 2.5-mL aliquot of the corresponding CO_2_ gas was injected through the septum. Chamber air was mixed by withdrawing and reinjecting 50 mL of air with a syringe ten times immediately after each gas addition. Plants remained in the chambers for a total of 3 h, after which they were induced by simulated herbivory. Volatiles were then collected on SuperQ filters for 3 h and analyzed by GC-MS. This analysis showed ^13^C incorporation into DMNT (Fig. S3A-C). No clear incorporation was found for the other induced volatiles.

Based on the preliminary results, receiver plants were pulse-labeled using the same procedure and placed individually in downstream chambers of the flow-through system. An unlabeled sender plant was placed in each upstream chamber, which was connected to its corresponding downstream receiver chamber. Sender plants were then induced by simulated herbivory, and emissions of labeled and unlabeled DMNT from sender plants alone and sender-receiver pairs were monitored by PTR-ToF-MS (Fig. 3D).

### Within-leaf volatile analysis

Within-leaf volatile pools were quantified using solid-phase microextraction followed by gas chromatography-mass spectrometry (SPME-GC-MS), as previously described (38). Fifty milligrams of frozen leaf powder were placed in a sealed 10 mL glass vial. A 100 μm polydimethylsiloxane-coated SPME fiber (Supelco, USA) was inserted and exposed to the headspace at 60°C for 35 min. Captured volatiles were thermally desorbed and analyzed using an Agilent 7820A gas chromatograph coupled to an Agilent 5977E quadrupole mass selective detector. Volatile compounds were identified by comparison with the NIST Mass Spectral Library (USA) and, when available, with authentic standards, and relative quantities were determined by integrating individual peak areas from extracted ion chromatograms (Fig. S9).

### Phytohormone analysis

The phytohormones 12-oxophytodienoic acid (OPDA), jasmonic acid (JA), JA-isoleucine (JA-Ile), abscisic acid (ABA), and salicylic acid (SA) were extracted from 100 mg of frozen leaf powder using ethyl acetate containing 1 ng of isotopically labeled internal standards (d_y_-JA, ¹³C₆-JA-Ile, d₆-ABA, d₆-SA) and analyzed by ultra-high-performance liquid chromatography coupled with tandem mass spectrometry (UHPLC-MS/MS), as previously described (76, 77).

### Gene expression analysis

Total RNA was extracted from 80 mg of frozen leaf tissue using the GeneJET Plant RNA Purification Kit (Thermo Scientific, USA). Genomic DNA was removed, and first-strand cDNA was synthesized using the PrimeScript RT Reagent Kit with gDNA Eraser (Takara Bio Inc., Japan). Quantitative reverse transcription PCR (qRT-PCR) was performed on a QuantStudio 5 Real-Time PCR System (Applied Biosystems) using ORA SEE qPCR Mix (highQu GmbH, Germany). Gene expression levels were normalized to *Actin*, and primer sequences are listed in Table S1.

### Stomatal conductance measurements

Stomatal conductance (gsw) was measured using a portable porometer (LI-600 Porometer/Fluorometer; LI-COR, Inc., Lincoln, NE, USA). Measurements were taken between the base and middle of each leaf at defined time points following treatment.

## Statistical analyses

All statistical analyses were performed in R version 4.4.2. For volatile emission time courses, receiver emissions were calculated as the signal measured from sender-receiver pairs minus the mean signal measured from sender plants alone at the same analyte and time point (Analyte × time). Emission time courses were then baseline-normalized by subtracting, for each analyte and treatment, the mean signal at time 0 h pooled across replicates, such that the group mean at 0 h equaled zero. Three parameters were quantified for each analyte from the baseline-normalized time courses: onset time, peak time, and total signal intensity. Onset time was estimated by change-point analysis using the cpt.meanvar function with the AMOC method implemented in the ‘changepoint’ package (78). Peak time was defined as the earliest time point at which the maximum signal occurred. Cumulative signal intensity was calculated as the trapezoidal area under the time course. Differences between sender and receiver plants were assessed using two-sided Wilcoxon rank-sum tests with Benjamini-Hochberg correction for multiple comparisons within each analyte. Internal volatile pools, transcript levels of volatile biosynthesis genes, and phytohormone concentrations were compared between sender and receiver plants at each time point using two-sided Wilcoxon rank-sum tests with Benjamini-Hochberg correction for multiple comparisons. Stomatal conductance data were analyzed using one-way analysis of variance (ANOVA). When ANOVA indicated significant treatment effects, pairwise comparisons were performed using Tukey’s honestly significant difference (HSD) test. Significance thresholds are indicated in figure legends.

## Acknowledgements

This work was supported by the Swiss National Science Foundation (Project Nr. 200355), the State Secretariat for Education, Research and Innovation (ERC CoG Replacement Scheme, SERI project No. MB22.00052), the National Natural Science Foundation of China (42377285), the Fundamental Research Funds for the Central Universities (226-2025-00049), and the University of Bern. We thank the research gardeners Christopher Ball and Sarah Dolder for plant cultivation, the laboratory of Prof. Jianqiang Wu for providing the *myc2ab* mutant, the laboratory of Prof. Anthony J. Studer for providing the *slac1* mutant, Prof. Michael Raissig and members of the Stomatal Biology Group at the University of Bern for advice and guidance on stomatal conductance measurements, and members of the Biotic Interactions and Chemical Ecology groups at the University of Bern for helpful discussions.

## Supplementary Figures

**Supplementary Figure S1.**
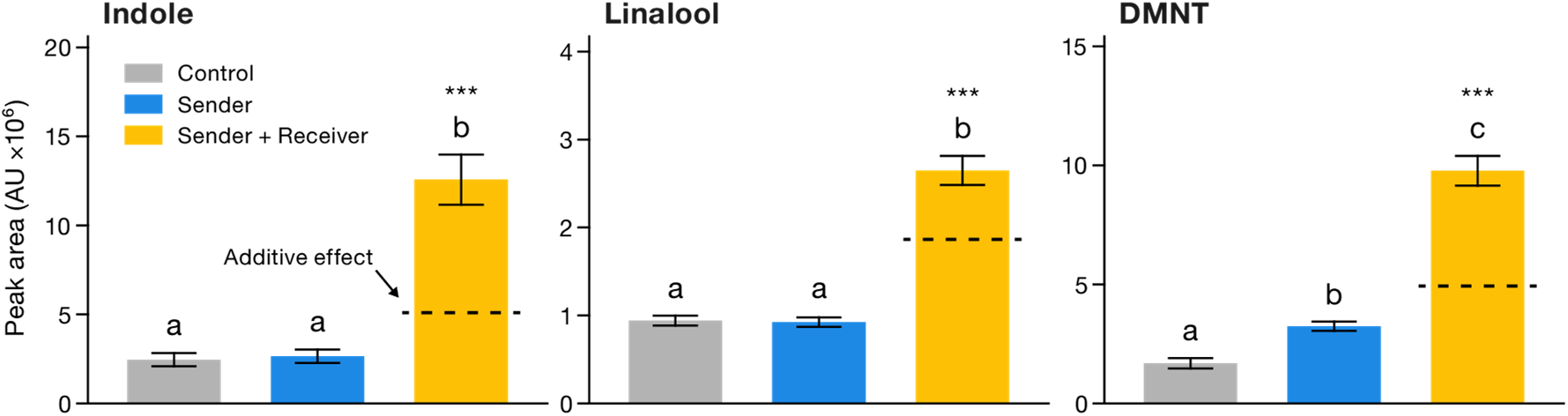
Early volatile emissions from sender-receiver pairs measured by GC-MS. Emissions of indole, linalool, and DMNT measured between 60–75 min after the first induction treatment from untreated control plants, herbivore-induced sender plants, and sender plants paired with undamaged receiver plants. Headspace volatiles were collected during this interval using SuperQ adsorbent traps and analyzed by gas chromatography-mass spectrometry (GC-MS). Data are presented as mean ± SE (n = 7). Different letters indicate significant differences among treatments, as determined by one-way ANOVA followed by Tukey’s honestly significant difference (HSD) test. Dashed lines represent the predicted additive emissions from herbivore-induced sender plants and untreated control plants. Asterisks indicate significant differences between sender–receiver pairs and the predicted additive values (*\*\*\*P* < 0.001).

**Supplementary Figure S2.**
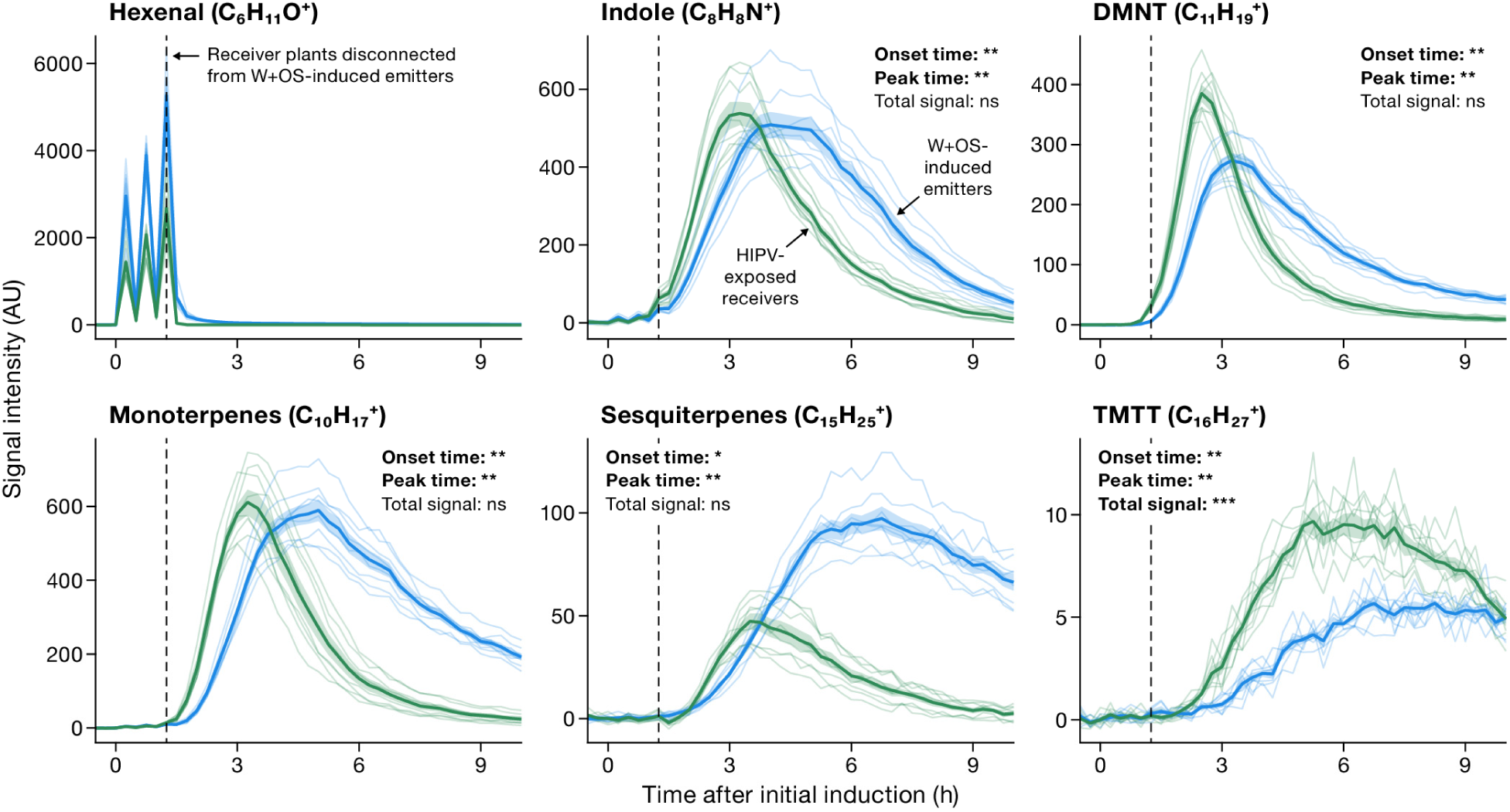
Earlier volatile emissions from receivers disconnected from senders after 75 min. Sender plants were induced by mechanical wounding followed by application of insect oral secretions at 0, 30, and 60 min. Receiver plants were exposed to sender emissions from 0 to 75 min and then disconnected from the senders (dashed line). Volatile emissions from W+OS-induced emitters (blue) HIPV-exposed, disconnected receivers (green) were monitored individually using PTR-ToF-MS. Thin lines show individual biological replicates; thick lines represent mean emission rates; shaded ribbons indicate ± SEM (n = 8). For each analyte, onset time, peak time, and cumulative signal intensity were quantified from baseline-corrected time courses as described for Figure 1. Sender and receiver values were compared using two-sided Wilcoxon rank-sum tests, with Benjamini-Hochberg correction across analytes within each metric. Asterisks indicate adjusted P values (P < 0.05; *P < 0.01; **P < 0.001); ns, not significant.

**Supplementary Figure S3.**
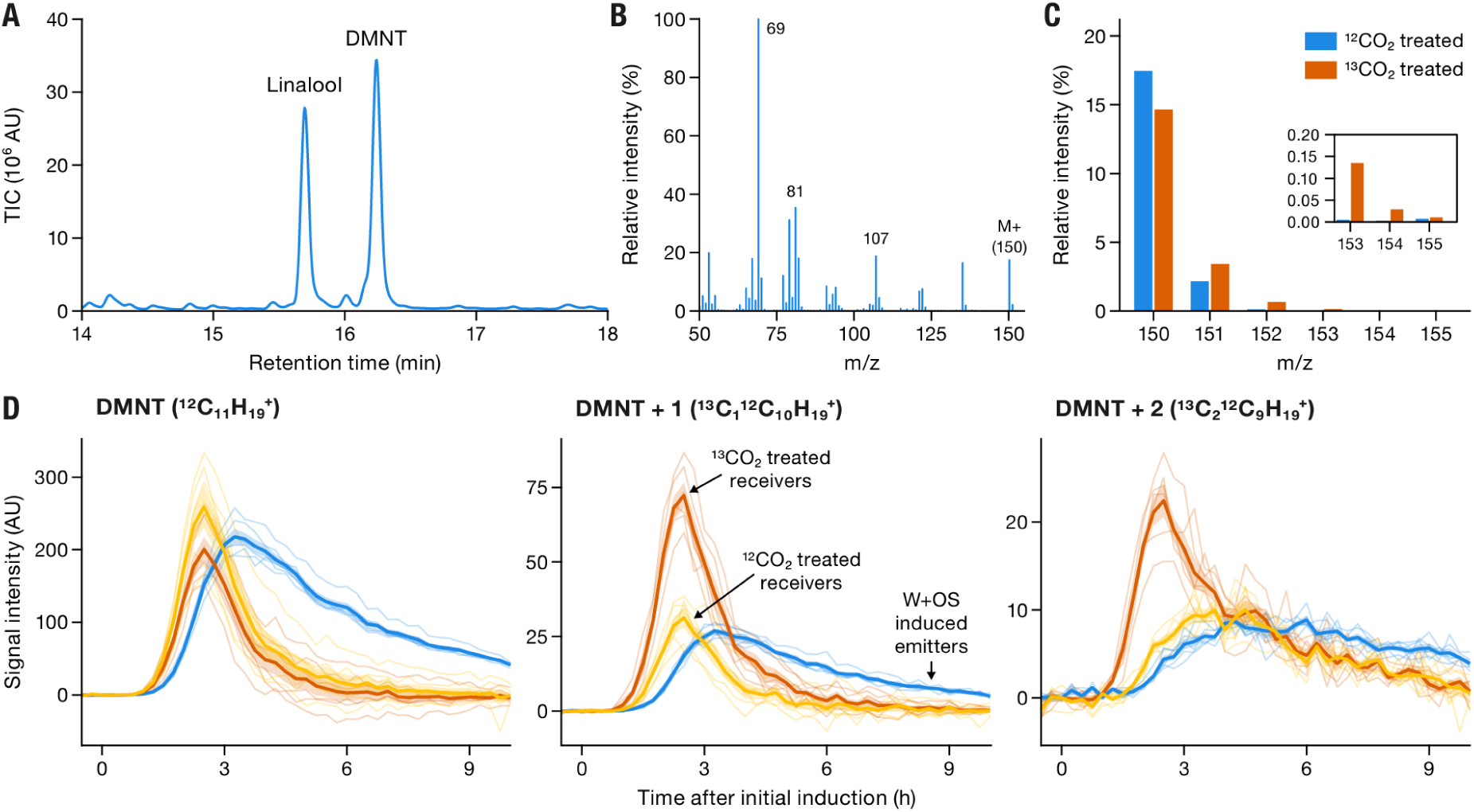
^13^CO_2_ labeling shows that receivers release *de novo* produced DMNT earlier than senders. (A) Representative GC-MS total ion chromatogram from a ^12^CO_2_-treated plant following W+OS induction, showing linalool and DMNT peaks. (B) Representative electron ionization mass spectrum of the DMNT peak. (C) Relative intensities of the molecular ion isotopologues of DMNT in W+OS-induced plants pretreated with ^12^CO_2_ or ^13^CO_2_; the inset enlarges the low-abundance m/z 153–155 ions. While significant incorporation of ^13^CO_2_ was detected for DMNT, no clear incorporation was found for linalool or other induced volatiles from the pulse labelling treatment. (D) Sender plants were induced by mechanical wounding followed by application of insect oral secretions at 0, 30, and 60 min and paired with receiver plants that had been pulse-labeled with ^13^CO_2_ (orange) or ^12^CO_2_ (yellow). Emissions of unlabeled DMNT and the +1 and +2 ^13^C isotopologues were monitored by PTR-ToF-MS. Sender emissions are shown in blue. Receiver emissions were calculated by subtracting mean sender emissions from emissions of sender-receiver pairs at each time point. Note that the later rising curves of the ^13^C_2_^12^C_9_H_19_^+^ signal (m/z 153.143) in the right panel are due to an insufficiently resolved, closely related m/z signal from another compound (m/z 153.128). Thin lines show individual biological replicates; thick lines represent mean emission rates; shaded ribbons indicate ± SEM (n = 6).

**Supplementary Figure S4.**
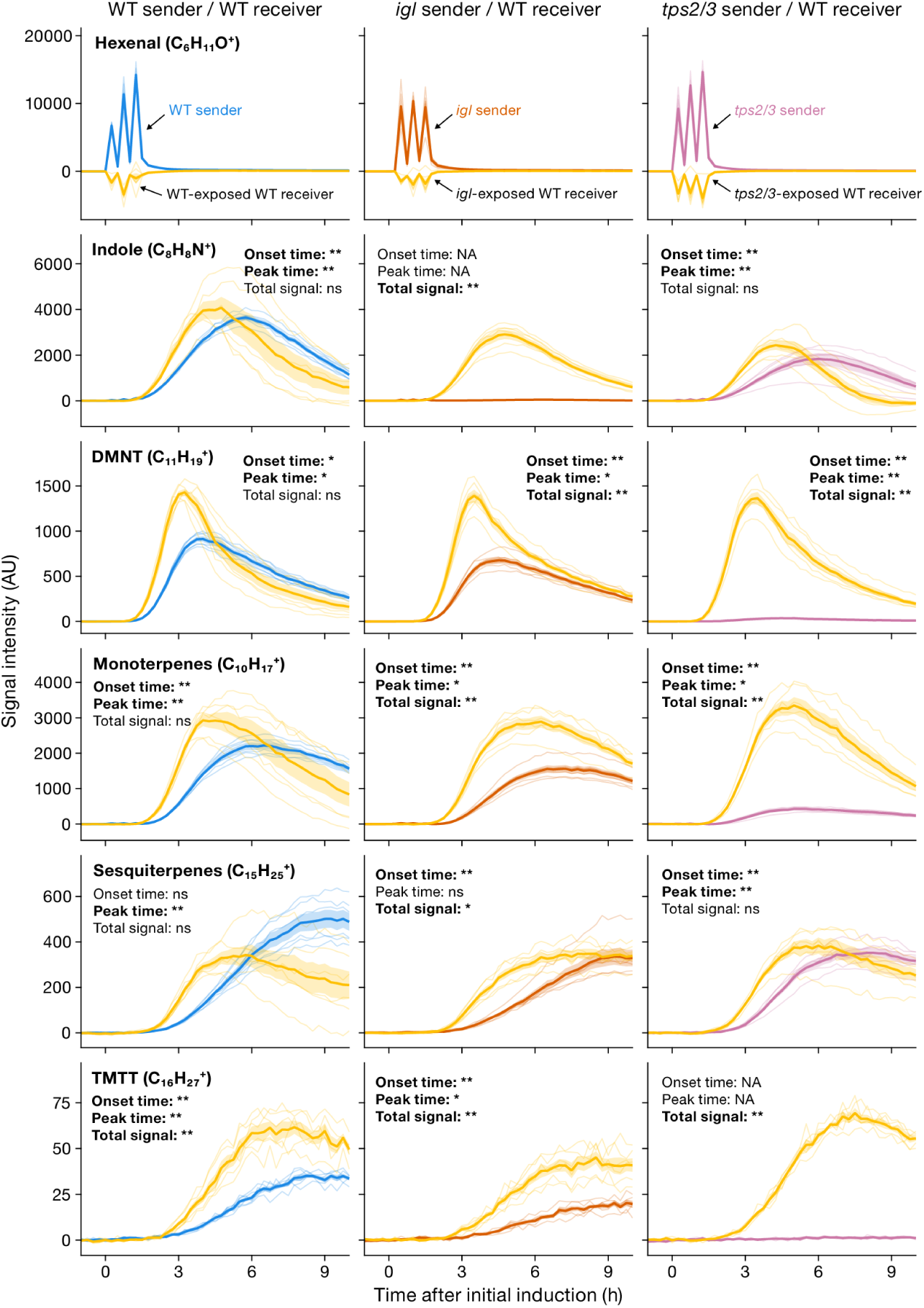
Indole, mono-and homoterpenes are not required for rapid responses in neighboring plants. Time-resolved volatile emissions were measured from wild-type (WT) receiver plants exposed to herbivory-induced WT or mutant sender plants carrying defects in major volatile biosynthesis pathways. Sender plants were induced by mechanical wounding followed by application of insect oral secretions at 0, 30, and 60 min, and volatile emissions were monitored every 15 min from sender plants alone and from sender-receiver pairs. Receiver emissions were calculated by subtracting emissions from sender plants alone from those measured in sender-receiver pairs. Shown are emissions from WT, *igl*, and *tps2/3* sender plants paired with WT receivers. The *igl* mutant is deficient in indole biosynthesis, whereas *tps2/3* is impaired in mono-and homoterpene biosynthesis. Stress-related volatiles, including (*Z*)-3-and (*E*)-2-hexenal (combined signal), indole, mono-, sesqui-, and homoterpenes (DMNT, TMTT), were detected as protonated ions based on characteristic *m/z* signals. Solid lines represent mean emission rates for sender plants (blue, orange, or pink) and WT receiver plants (yellow); shaded ribbons indicate ± SEM, and thin lines show individual biological replicates (n = 6). For each analyte, onset time, peak time, and cumulative signal intensity were quantified from baseline-normalized time courses. Statistical differences were assessed using two-sided Wilcoxon rank-sum tests with Benjamini-Hochberg correction for multiple comparisons. Asterisks indicate significance levels (\**P* < 0.05; \*\**P* < 0.01); ns, not significant; NA, metric not determined. For other mutants, see Fig. 4.

**Supplementary Figure S5.**
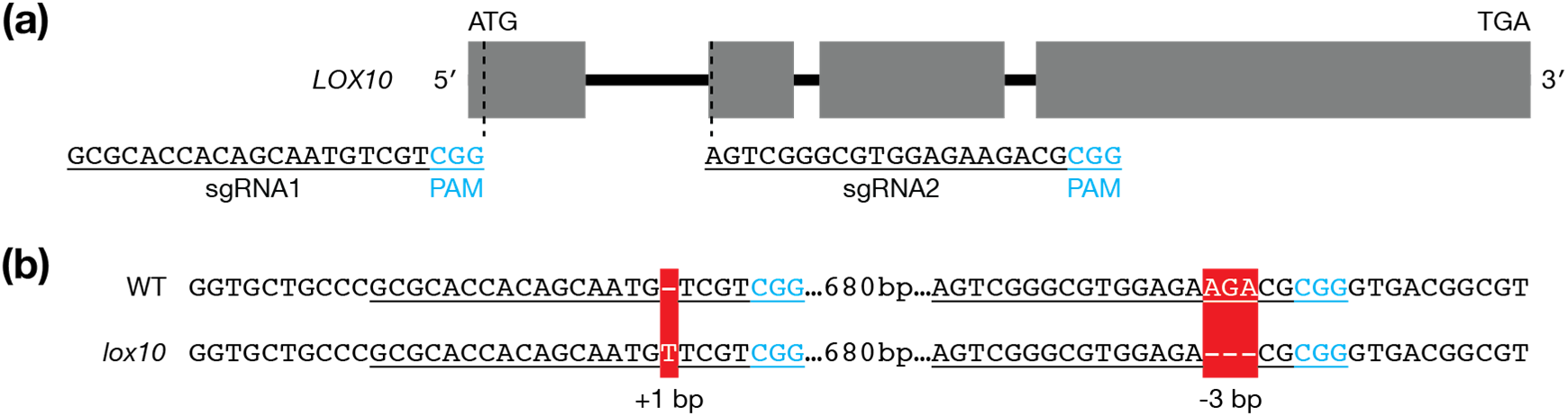
(a) Schematic representation of the *LOX10* gene structure showing exon-intron organization (gray boxes, exons; black lines, introns). Two single guide RNAs (sgRNA1 and sgRNA2) were designed to target the 5′ region of the coding sequence to maximize the likelihood of generating a loss-of-function allele. The protospacer-adjacent motif (PAM) sequences are indicated in blue. (b) Sequence alignment of the targeted regions in wild type (WT) and the *lox10* mutant allele. Red boxes indicate mutations at the sgRNA target sites relative to WT. A one-base insertion (+1 bp) was detected at the first target site (sgRNA1), and a three-base deletion (−3 bp) was identified at the second target site (sgRNA2). PAM sequences are highlighted in blue and target sequences are underlined. These mutations are predicted to disrupt *LOX10* function.

**Supplementary Figure S6.**
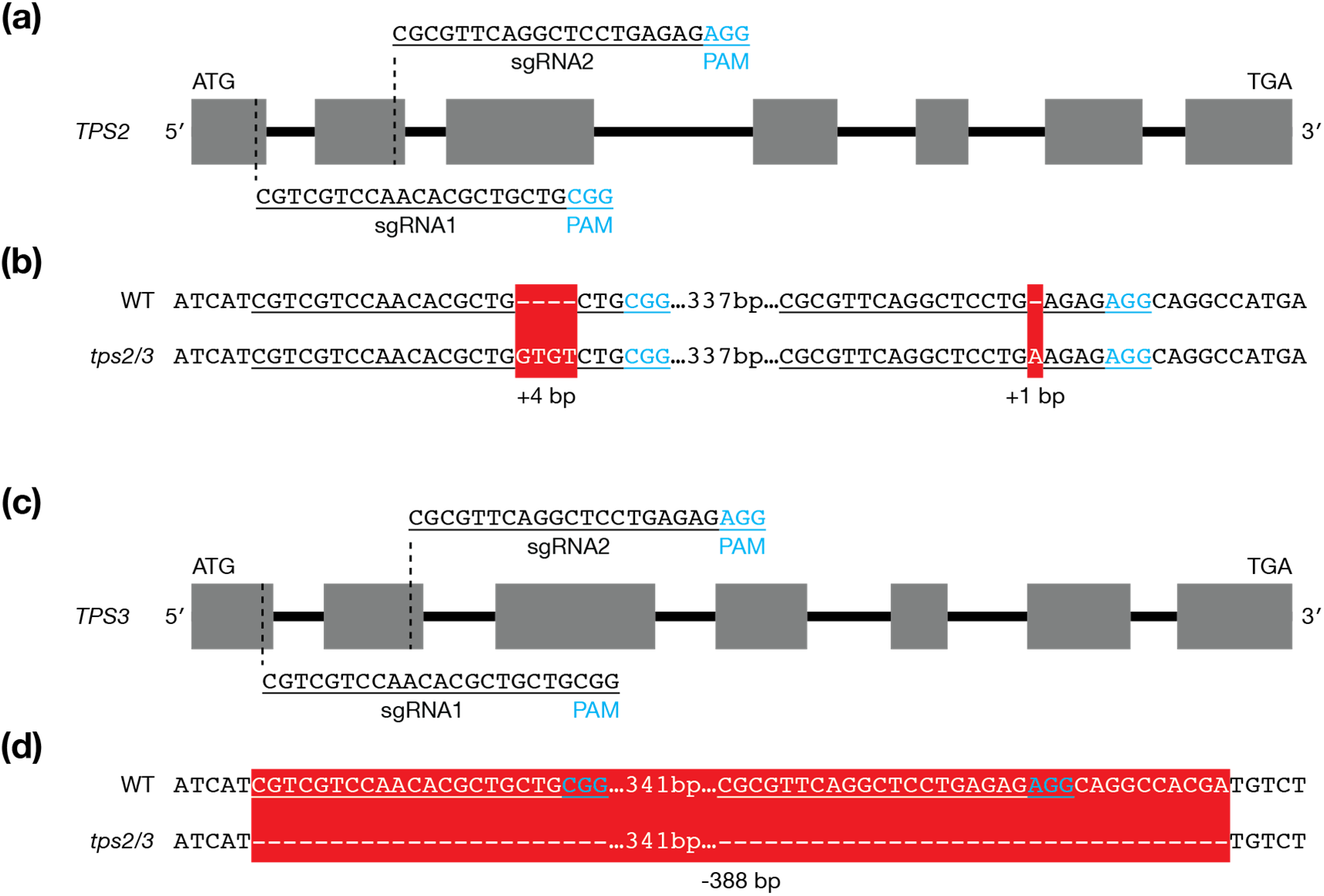
(a) Schematic representation of the *TPS2* gene structure showing exon-intron organization (gray boxes, exons; black lines, introns). Two single guide RNAs (sgRNA1 and sgRNA2) targeting conserved coding regions near the 5′ end were designed. The protospacer adjacent motif (PAM) sequences are indicated in blue. (b) Sequence alignment of the targeted regions in wild type (WT) and the *tps2/3* mutant for *TPS2*. Red boxes indicate insertions at the sgRNA1 (+4 bp) and sgRNA2 (+1 bp) target sites relative to WT. PAM sequences are highlighted in blue and target sequences are underlined. The mutations result in frameshifts predicted to disrupt *TPS2* function. (c) Schematic representation of the *TPS3* gene structure with the same sgRNA1 and sgRNA2 target sites as used for *TPS2*. Exons and introns are indicated as in (a), and PAM sequences are shown in blue. (d) Sequence alignment of the targeted regions in WT and *tps2/3* for *TPS3*. A large deletion of 388 bp between the two sgRNA target sites was detected in the mutant allele. The deleted region (red box) spans both target sites, leading to a frameshift and predicted loss of *TPS3* function.

**Supplementary Figure S7.**
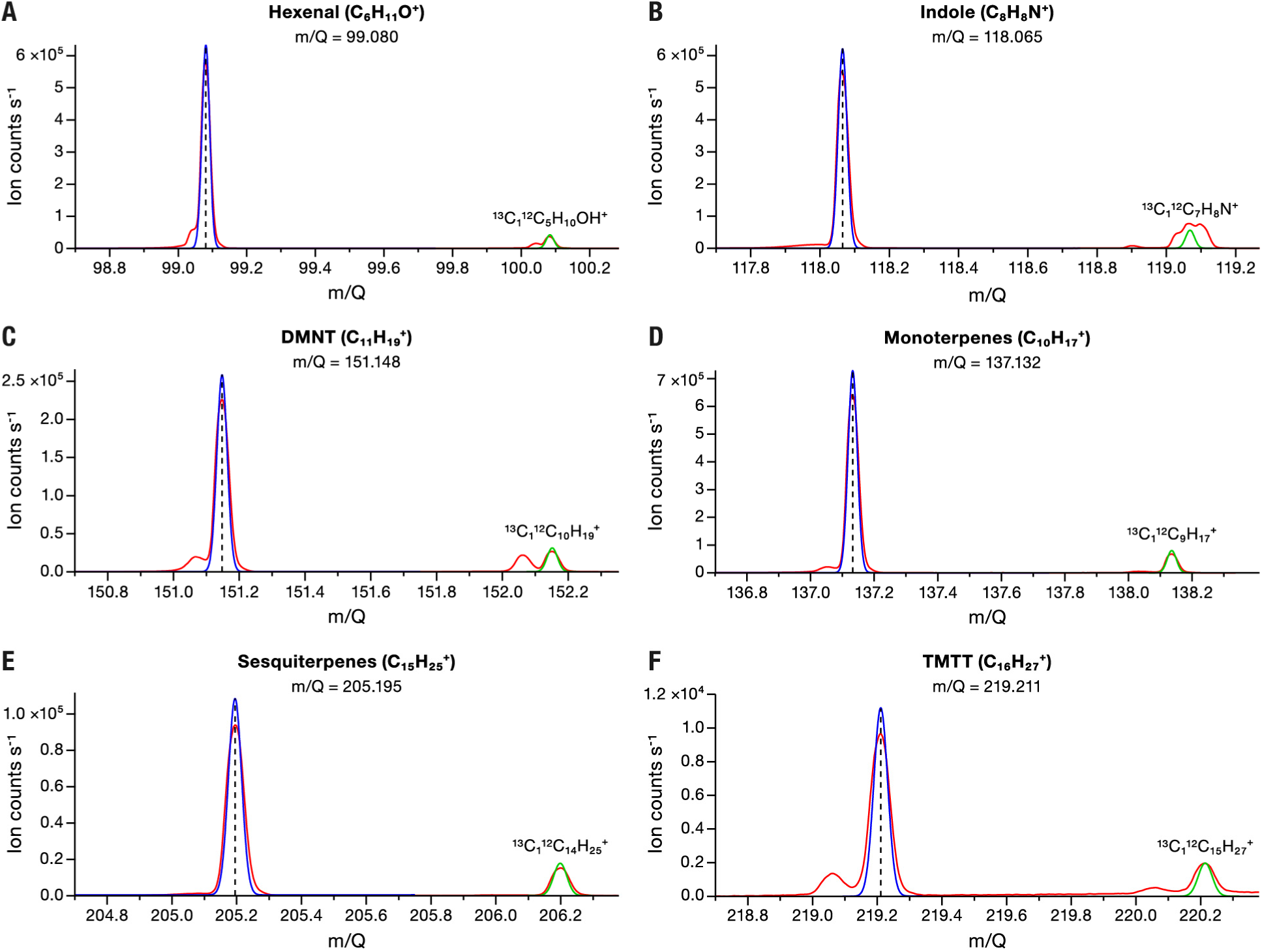
High-resolution PTR-ToF-MS spectra support assignment of monitored volatile ions. Mass spectra are shown for (A) hexenal, (B) indole, (C) DMNT, (D) monoterpenes, (E) sesquiterpenes, and (F) TMTT. The expected monoisotopic ion is marked by a dashed line, and the corresponding ^13^C isotopologue is shown at the expected higher m/Q.

**Supplementary Figure S8.**
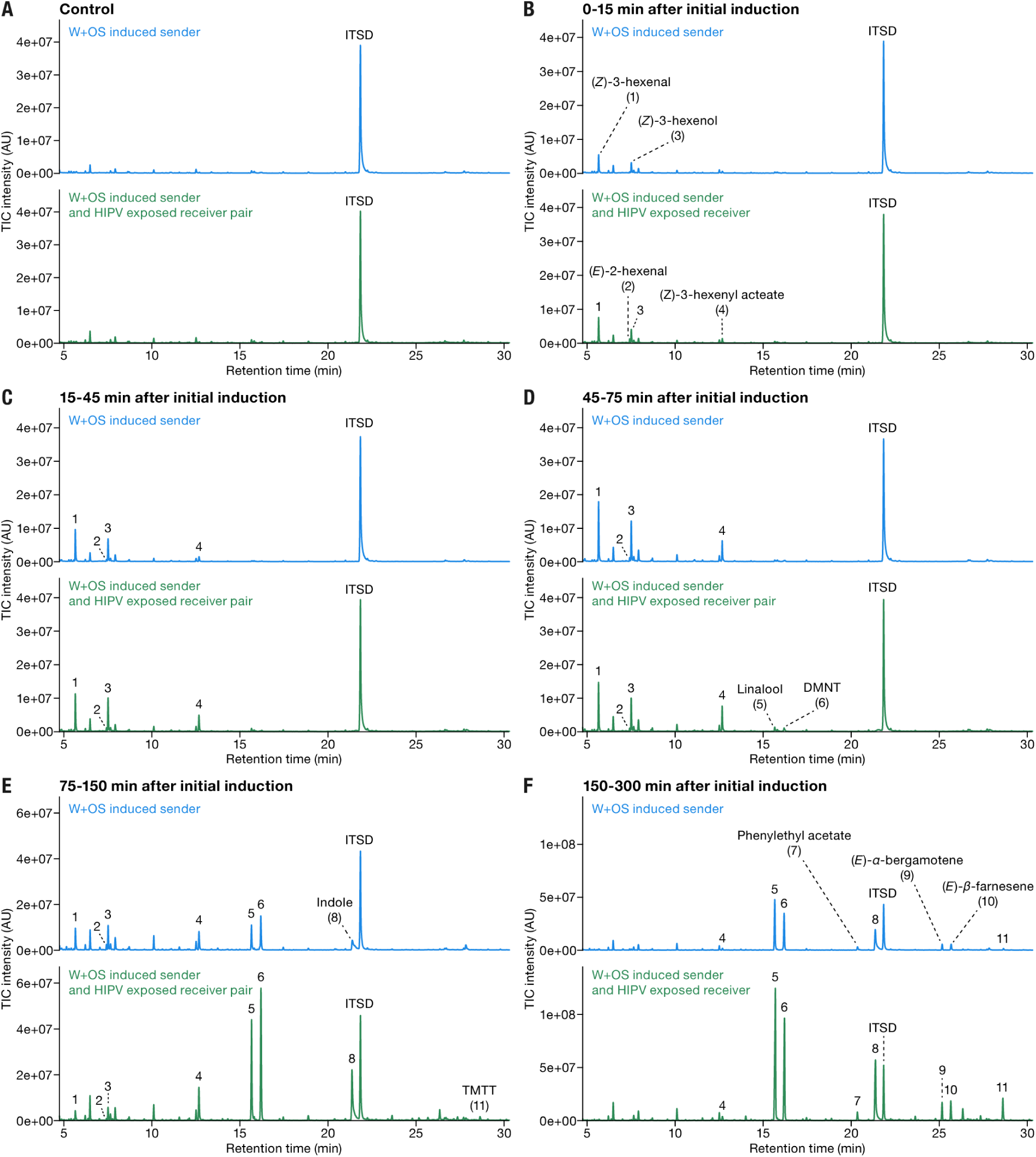
GC-MS chromatograms of volatile emissions from W+OS-induced sender plants and sender-receiver pairs. Sender plants were induced by mechanical wounding followed by application of insect oral secretions at 0, 30, and 60 min. Representative total ion chromatograms of SuperQ-collected headspace samples are shown for untreated controls (A) and samples collected 0–15 min (B), 15–45 min (C), 45–75 min (D), 75–150 min (E), and 150–300 min (F) after initial induction. Sender plants alone are shown in blue, and sender plants paired with HIPV-exposed receivers are shown in green. Identified peaks are (1) (*Z*)-3-hexenal, (2) (*E*)-2-hexenal, (3) (*Z*)-3-hexenol, (4) (*Z*)-3-hexenyl acetate, (5) linalool, (6) DMNT, (7) phenylethyl acetate, (8) indole, (9) (*E*)-*α*-bergamotene, (10) (*E*)-*β*-farnesene, and (11) TMTT. ITSD, nonyl acetate internal standard.

**Supplementary Figure S9.**
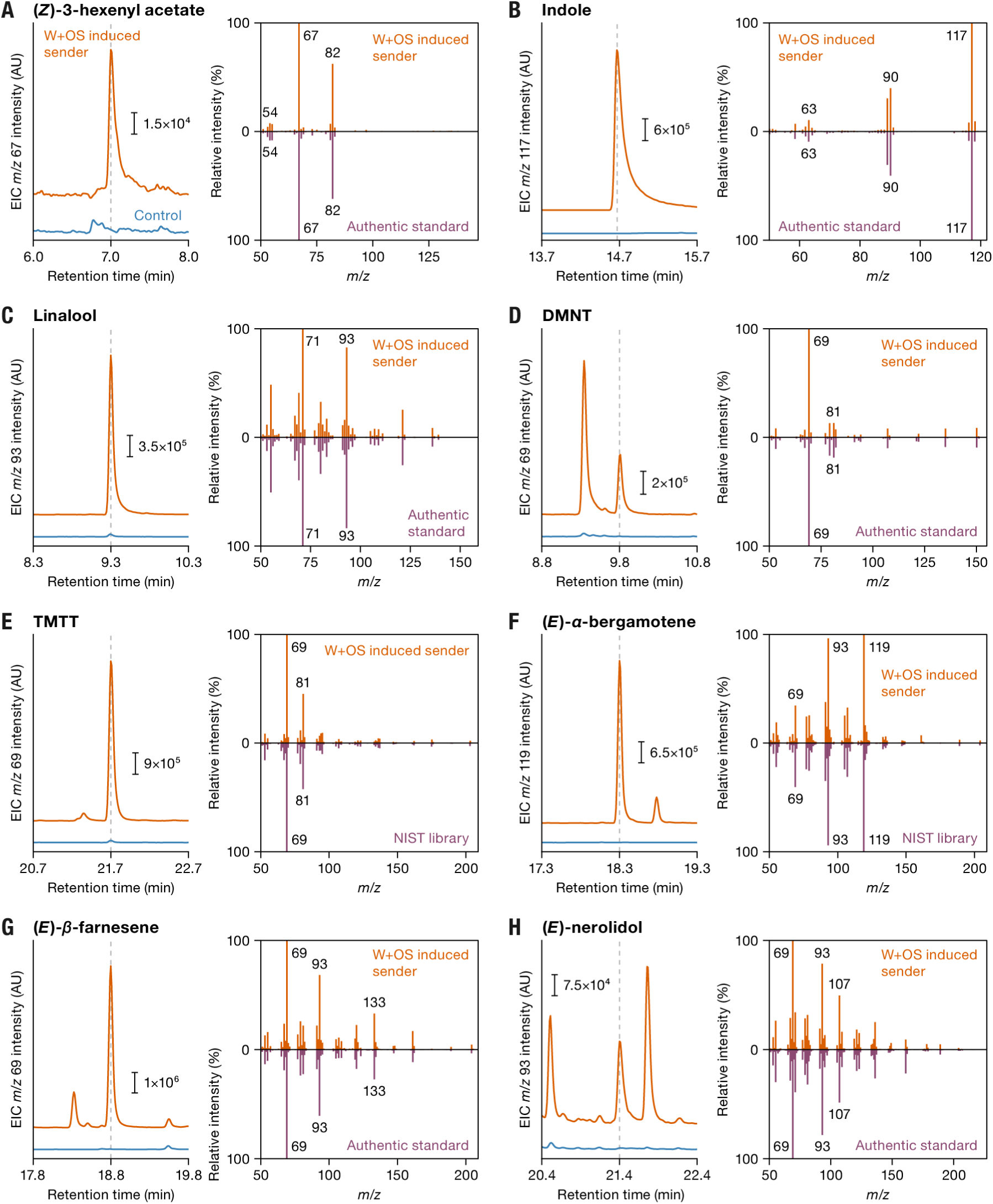
SPME-GC-MS validation of within-leaf volatile compound assignments. Representative extracted ion chromatograms from untreated control leaves (blue) and W+OS-induced sender leaves (orange), together with their corresponding electron ionization mass spectra. Mirrored spectra show authentic standards (purple) for (A) (*Z*)-3-hexenyl acetate, (B) indole, (C) linalool, (D) DMNT, (G) (*E*)-*β*-farnesene, and (H) (*E*)-nerolidol, and NIST library spectra for (E) TMTT and (F) (*E*)-*α*-bergamotene. Dashed lines indicate the retention time of each assigned compound.

## Supplementary Tables

**Supplementary Table S1.**
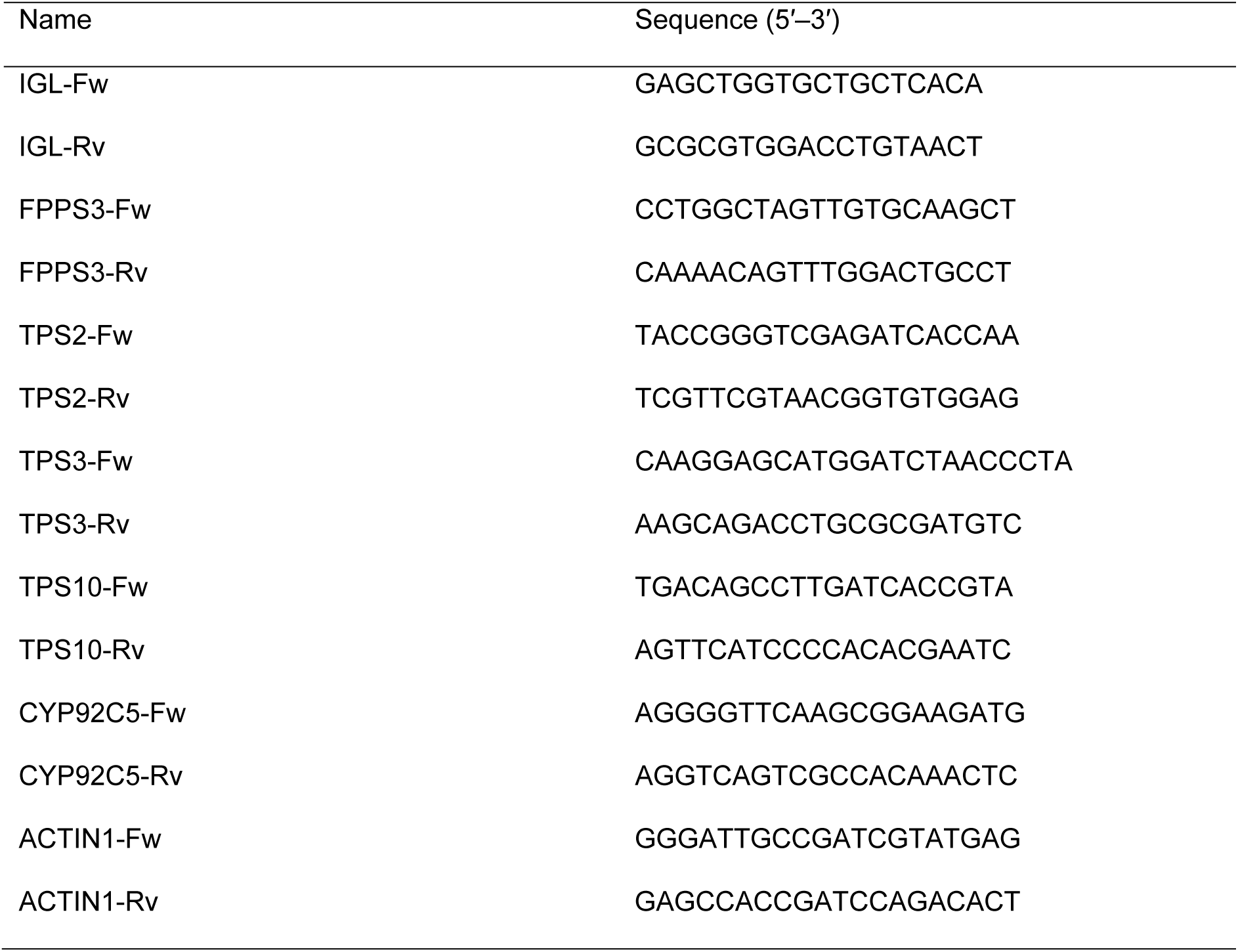
Primers used in this study.

